# Complete mitogenome of Kashmir musk deer (*Moschus cupreus*) and its comparative phylogenetic relationships

**DOI:** 10.1101/2020.08.25.265850

**Authors:** Bhim Singh, Kumudani Bala Gautam, Subhashree Sahoo, Ajit Kumar, Sandeep Kumar Gupta

**Author notes:** Address for Correspondence, **Dr. S. K. Gupta**, Scientist-E, Wildlife Institute of India, Chandrabani, Dehra Dun, 248 001 (U.K.), India.

## Abstract

The endangered Kashmir musk deer (*Moschus cupreus*) is native to the high altitudinal region of the Himalayas. In this study, we sequenced, annotated and characterized the complete mitogenome of *M. cupreus* to gain insight into the molecular phylogeny and evolution of musk deer. The mitogenome of *M. cupreus*, which is 16,354 bp long comprised 13 protein-coding genes (PCGs), 22 transfer RNA genes (tRNAs), two ribosomal RNA genes (rRNAs) and non-coding control region. The *M. cupreus* mitogenome composition was highly A+T biased 68.42%, and exhibited a positive AT skew (0.082) and negative GC skew (0.307). The phylogenetic analysis suggested that KMD is the most primitive extant species in the genus *Moschus* whereas Alpine musk deer (*M. chrysogaster*) and Himalayan musk deer (*M. leucogaster*) are closely related. This result confirmed the placement of *M. cupreus* within the monotypic family Moschidae of musk deer. This study provides a better understanding of lineage identification and musk deer evolution for further research.

## Introduction

The Kashmir musk deer (*Moschus cupreus*) is a member of the genus *Moschus* in the monotypic family Moschidae (Grubb 1982). The species are endemic to the Palearctic region and mostly inhabit high altitudinal mountainous regions of the Himalayan ranges (Grubb 1982). According to the IUCN database, the current population is in decline due to overexploitation, primarily caused by habitat destruction and illegal hunting of musk pod and skin in wildlife trade. The IUCN Red List includes *M. cupreus* in the ‘Endangered’ category and has listed in Appendix I under the Convention on International Trade in Endangered Species of Wild Fauna and Flora (CITES). In India, musk deer is categorized under the Schedule I of the Indian Wildlife Protection Act, 1972. Due to limited ecological and genetic evidence, the distribution range and evolutionary history of the species are still unclear (Pan et al., 2015). According to Grubb 2005, the distribution of *M. cupreus* is restricted to the Himalayan region of Kashmir, Pakistan and Afghanistan. However, a recent molecular study has provided a new distribution record of *M. cupreus* from Mustang, Nepal and west of Annapurna Himalayas range (Singh et al., 2019). The availability of complete mitogenomes will furnish molecular evidence that will resolve distribution, evolutionary as well as phylogenetic dilemmas. Out of the seven *Moschus* species, mitogenomes of *M. cupreus* and *M. fuscus* are still lacking. Due to maternal mode of inheritance, presence of conserved genes, rare recombination and high evolutionary rate, the mitogenomic data has been widely employed in phylogenetics and evolutionary genomics (Avise et al., 1985, Boore 1999). Hence, it warrants a comprehensive phylogenetic study of *M. cupreus* for clear species delimitation which would help in designing scientific strategies for conservation breeding musk deer.

Therefore, we sequenced and characterized for the first time, the complete mitogenome of *M. cupreus*. We further investigated the phylogenetic relationship of *M. cupreus* with other *Moschus* species.

## Materials and methods

Two tissue samples of *M. cupreus* were used from the repository of Wildlife Forensic and Conservation Genetics (WFCG) Cell of our institute. These samples were forwarded by the State Forest Departments of Jammu & Kashmir and Uttarakhand. Genomic DNA (gDNA) was extracted from the samples using the DNeasy Blood Tissue Kit (QIAGEN, Germany) in a final elution volume of 100 μl. The extracted DNA was checked by 0.8% agarose gel and thereafter diluted in final concentration in 40 ng/μl for further PCR amplification.

### PCR amplification and sequencing

The complete mtDNA genome was amplified and sequenced by using 21 different sets of primers (Hassanin et al., 2009). In addition to the available set of primers, 3 fragments did not successfully amplify, therefore, we designed novel primers to cover the gaps (Supplementary file: ST1). DNA amplifications were performed in 20 μl reaction volume containing 1 × PCR buffer (50mM KCl, 10mM Tris–HCl, and pH 8.3), 1.5mM MgCl2, 0.2mM of each dNTPs, 3 pmol of each primer, 5U of DreamTaq DNA Polymerase and 1 μl (~30 ng) of template DNA. The PCR protocol was set as follows: initial denaturation at 95 °C for 10 min, followed by 35 cycles of denaturation at 95 °C for 45 s, annealing at 55 °C for 40 s and extension at 72 °C for 75 s. The final extension was undertaken at 72 °C for 10 min. PCR amplification was confirmed by electrophoresis and visualized under UV transilluminator. To remove unwanted residual primers, the PCR products were then treated with Exonuclease-I and Shrimp Alkaline Phosphatase (Thermo Scientific Inc.) at 37°C for 20 min and at 85°C for 15 min. The purified fragments were sequenced directly in an Applied Biosystems Genetic Analyzer, ABI 3500 XL in both directions.

### Mitogenome alignment and annotation

The complete mitogenome of *M. cupreus* was generated by aligning the overlapping fragments of DNA sequences using Sequencher^®^ version 5.4.6 (Gene Codes Corporation, Ann Arbor, MI, USA). Annotation of mitogenome was performed by using the Mitos Web Server (Bernt et al., 2013). The complete mtDNA of 16,354 bp was obtained and the gene map of *M. cupreus* was generated using CGView Server (Grant et al., 2008). MEGA X (Kumar et al., 2018) was implemented to calculate the base composition in the mtDNA genetic code. Bias in nucleotide composition among the complete mitogenome of *M. cupreus* was estimated using skew analysis where: AT skew = (A-T)/(A+T) and GC skew = (G-C)/(G+C) (Perna et al., 1995). The Relative Synonymous Codon Usage (RSCU) values of 13 protein-coding genes (PCGs) of *M. cupreus* were calculated using MEGA X (Kumar et al., 2018). The intergenic spacer and overlapping regions interspersed between genes of complete mitogenome were estimated manually.

### Phylogenetic analysis and genetic differentiation

The phylogenetic analysis was performed using the whole mitogenome of *M. cupreus* sequenced along with available GenBank database of *Moschus* species: Alpine musk deer (*M. chrysogaster* KP684123 and KC425457), Himalayan musk deer (*M. leucogaster* NC042604 and MG602790), Siberian musk deer (*M. moschiferus* FJ469675), Anhui musk deer (*M. anhuiensis* KP684124 and KC013352) and Forest musk deer (*M. berezovskii* EU043465). To get a better insight into the phylogeny of musk deer, we also included eight mitogenomes representing Cervini, Bovini, and Boselaphini from GenBank. Besides, one sequence of *Moschiola indica* (KY290452) was chosen as an outgroup. A bayesian consensus tree was constructed in BEAST version 1.7 (Drummond et al., 2012). Bayesian inference analysis was finalized by using Monte Carlo Markov Chain (MCMC) chain method for 10 million generations and sampling one tree at every 1000 generations using a burn-in of 5000 generations. FigTree v1.4.0 (http://tree.bio.ed.ac.uk/software/figtree/) was used to visualize the phylogenetic tree. We estimated the similarity of complete mitogenomes among musk deer species using MatGAT (Campanella et al., 2003). Additionally, we calculated gene-specific mean pairwise genetic distance within the musk deer species to elucidate differentiation among the 13 PCGs using MEGAX (Kumar et al., 2018) implementing Tamura 3-Parameter Model (T92 + G).

## Results and discussion

### Mitogenome organization

We obtained a 16,354 bp long circular mitochondrial genome of Kashmir Musk deer (*M. cupreus*) which has been submitted in the GenBank (MTXXXXXX and MTXXXXXX). It showed the typical mammalian mitogenomic arrangement with 22 tRNA genes, 13 protein-coding genes (PCGs), two rRNA genes and a non-coding hypervariable control region (Fig. 1). The mitogenome arrangement and gene distribution of *M. cupreus* showed similarity with other *Moschus* species (Jang et al., 2010; Guo et al., 2018). The total nucleotide composition in *M. cupreus* was: A (33.9%), T (28.8%), C (24.4%) and G (12.9%) (Table 2). Apart from ND6 gene, the other eight tRNA genes (tRNAGln, tRNAAla, tRNAAsn, tRNACys, tRNATyr, tRNASer, tRNAGlu, tRNAPro) were also encoded on the light strand (L-strand) whereas remaining genes were encoded on the heavy strand (H-strand). Six pairs of overlapping regions in the mitogenome were observed among tRNA-Val / 16S rRNA, tRNA-Ile / tRNA-Gln, COI / tRNA-Ser, ATP8 / ATP6, ND4L / ND4 and tRNA-Thr / tRNA-Pro.

**Figure 1:**
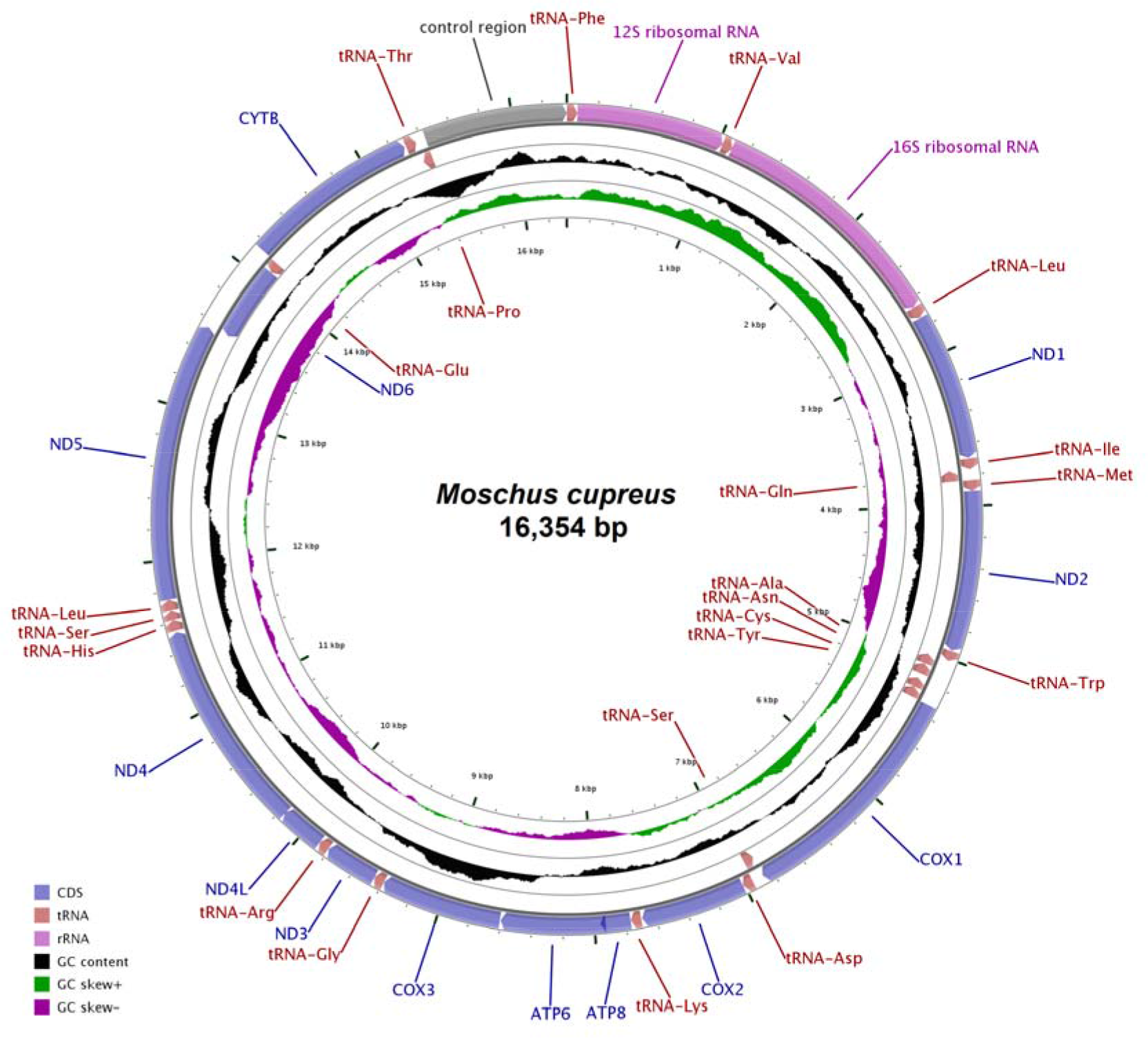
Illustration of the location of genes on the complete mitochondrial genome of *M.cupreus*.

**Table 1:**
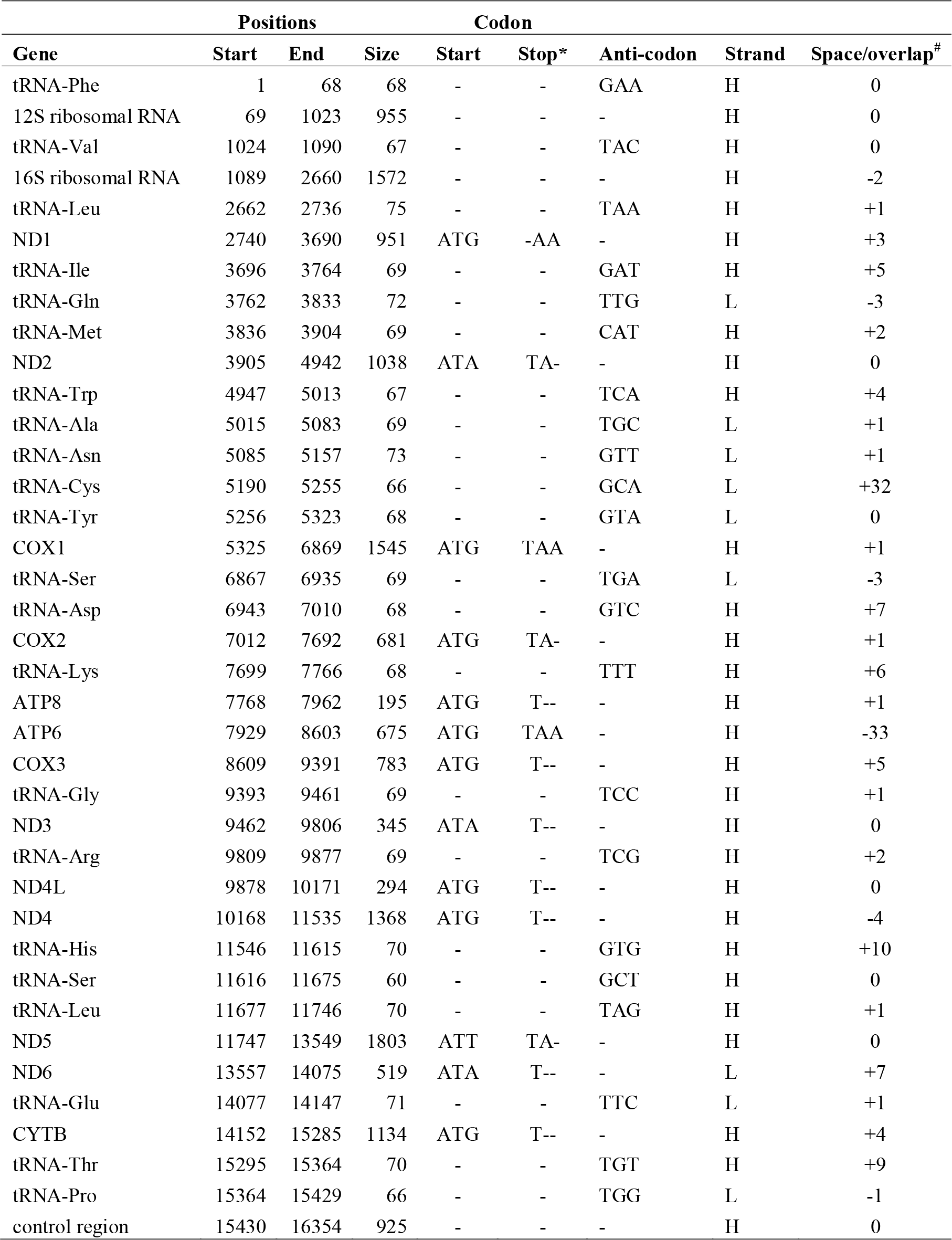
Organization of the mitochondrial genome of *M. cupreus*.

| Gene | Positions |  |  | Codon |  |  | Strand | Space/overlap<sup>#</sup> |
| --- | --- | --- | --- | --- | --- | --- | --- | --- |
|  | Start | End | Size | Start | Stop* | Anti-codon |  |  |
| tRNA-Phe | 1 | 68 | 68 | - | - | GAA | H | 0 |
| 12S ribosomal RNA | 69 | 1023 | 955 | - | - | - | H | 0 |
| tRNA-Val | 1024 | 1090 | 67 | - | - | TAC | H | 0 |
| 16S ribosomal RNA | 1089 | 2660 | 1572 | - | - | - | H | -2 |
| tRNA-Leu | 2662 | 2736 | 75 | - | - | TAA | H | +1 |
| ND1 | 2740 | 3690 | 951 | ATG | -AA | - | H | +3 |
| tRNA-Ile | 3696 | 3764 | 69 | - | - | GAT | H | +5 |
| tRNA-Gln | 3762 | 3833 | 72 | - | - | TTG | L | -3 |
| tRNA-Met | 3836 | 3904 | 69 | - | - | CAT | H | +2 |
| ND2 | 3905 | 4942 | 1038 | ATA | TA- | - | H | 0 |
| tRNA-Trp | 4947 | 5013 | 67 | - | - | TCA | H | +4 |
| tRNA-Ala | 5015 | 5083 | 69 | - | - | TGC | L | +1 |
| tRNA-Asn | 5085 | 5157 | 73 | - | - | GTT | L | +1 |
| tRNA-Cys | 5190 | 5255 | 66 | - | - | GCA | L | +32 |
| tRNA-Tyr | 5256 | 5323 | 68 | - | - | GTA | L | 0 |
| COX1 | 5325 | 6869 | 1545 | ATG | TAA | - | H | +1 |
| tRNA-Ser | 6867 | 6935 | 69 | - | - | TGA | L | -3 |
| tRNA-Asp | 6943 | 7010 | 68 | - | - | GTC | H | +7 |
| COX2 | 7012 | 7692 | 681 | ATG | TA- | - | H | +1 |
| tRNA-Lys | 7699 | 7766 | 68 | - | - | TTT | H | +6 |
| ATP8 | 7768 | 7962 | 195 | ATG | T-- | - | H | +1 |
| ATP6 | 7929 | 8603 | 675 | ATG | TAA | - | H | -33 |
| COX3 | 8609 | 9391 | 783 | ATG | T-- | - | H | +5 |
| tRNA-Gly | 9393 | 9461 | 69 | - | - | TCC | H | +1 |
| ND3 | 9462 | 9806 | 345 | ATA | T-- | - | H | 0 |
| tRNA-Arg | 9809 | 9877 | 69 | - | - | TCG | H | +2 |
| ND4L | 9878 | 10171 | 294 | ATG | T-- | - | H | 0 |
| ND4 | 10168 | 11535 | 1368 | ATG | T-- | - | H | -4 |
| tRNA-His | 11546 | 11615 | 70 | - | - | GTG | H | +10 |
| tRNA-Ser | 11616 | 11675 | 60 | - | - | GCT | H | 0 |
| tRNA-Leu | 11677 | 11746 | 70 | - | - | TAG | H | +1 |
| ND5 | 11747 | 13549 | 1803 | ATT | TA- | - | H | 0 |
| ND6 | 13557 | 14075 | 519 | ATA | T-- | - | L | +7 |
| tRNA-Glu | 14077 | 14147 | 71 | - | - | TTC | L | +1 |
| CYTB | 14152 | 15285 | 1134 | ATG | T-- | - | H | +4 |
| tRNA-Thr | 15295 | 15364 | 70 | - | - | TGT | H | +9 |
| tRNA-Pro | 15364 | 15429 | 66 | - | - | TGG | L | -1 |
| control region | 15430 | 16354 | 925 | - | - | - | H | 0 |

**Table 2:**
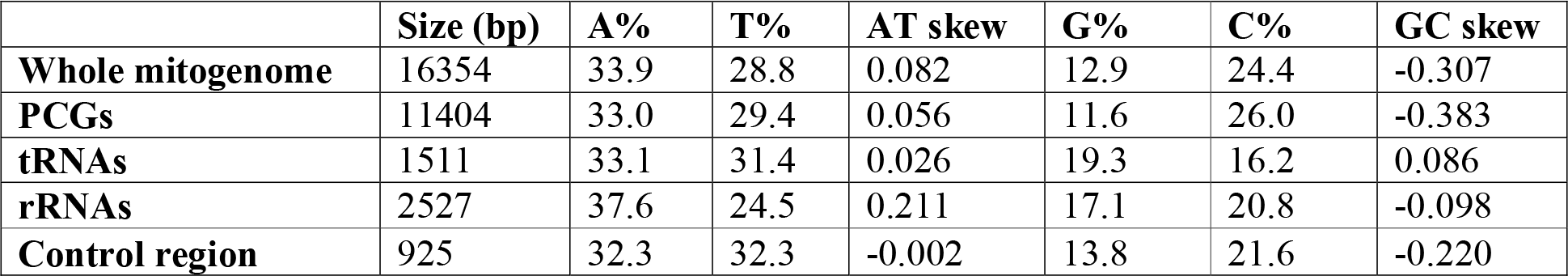
Nucleotide composition and Skewness in the complete mitogenome of *M. cupreus*.

The overlapping region ranges from 1 to 33 bp. The smallest overlapping region was found between tRNA-Thr / tRNA-Pro (−1 bp) whereas the longest was between ATP8 / ATP6 (−33 bp). Between tRNA-Pro and tRNA-Phe, a hypervariable control region was positioned. Furthermore, 22 intergenic spacers were found in between the mitochondrial regions which range from 1 to 32 bp length, while the longest space was found between tRNA-Asn and tRNA-Cys (Table 1). To identify nucleotide composition in the complete mitogenome of *M. cupreus*, we examined the values of AT-skew, GC-skew, and AT%. The resultant value of AT-skew was positive whereas the value of GC-skew was negative. The A+T and G+C content in the whole mitogenome was found to be AT biased with 62.5% and 37.5% respectively (Table 2).

### Protein coding genes (PCGs)

The total length of the 13 PCGs was estimated at 11,404 bp in *M. cupreus* mitogenome that included 37 bp overlapping fragments constituting 69.50% of the mitogenome (Table 2). The nucleotide composition of PCGs was A = 33% T = 29.5%, G = 11.6 and C=26. We found a higher abundance of AT% than GC%. The *M. cupreus* PCGs comprised twelve majority strand or H-strand genes (NADH dehydrogenases: ND1, ND2, ND3, ND4, ND5, and ND4L; three cytochrome c oxidases: COI, COII, and COIII; two ATPases: ATP6 and ATP8, and one cytochrome b: Cyt *b* gene) and one minority strand or L-strand gene (NADH dehydrogenase: ND6 gene) (Table 1). To understand the nucleotide distribution in PCGs, we estimated base skewness resulting in AT and GC skew of 0.056 and −0.383 receptively in PCGs. This suggests that in *M. cupreus*, an abundance of A is significantly higher than that of T while the abundance of C is significantly higher than that of G (Table 2). All the 13 PCGs started with ATN codon (ATG or ATA or ATT), congruent with other musk deer species. Excluding the stop codons, the relative synonymous codon usage (RSCU) for the 13 PCGs of *M. cupreus* comprised 3801 codons (Fig. 2).

**Figure 2:**
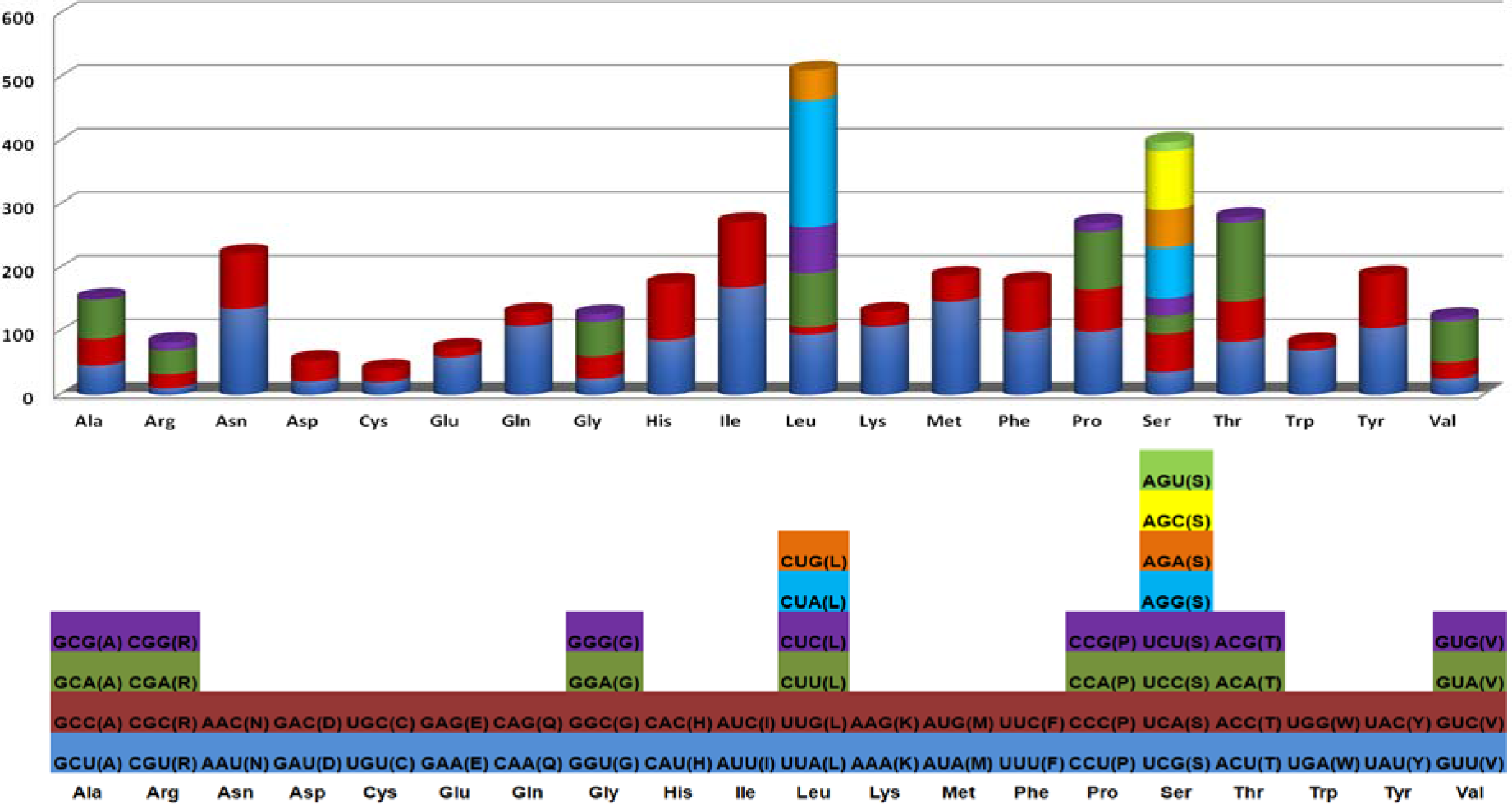
Relative synonymous codon usage (RSCU) of the mitochondrial protein-coding genes of the *M. cupreus* mitochondrial genome. Codon count numbers are provided on the X-axis.

### Ribosomal RNA and transfer RNA genes

In the *M. cupreus* mitogenome, two ribosomal RNA and twenty-two tRNA genes were recognized. The 12S rRNA was 955 bp long while 16S rRNA was 1572 bp long. The 12S rRNA gene was situated between tRNA-Phe and tRNA-Val, and likewise, the 16S rRNA gene was situated between tRNA-Val and tRNA-Leu. The total AT content of the two ribosomal rRNA was 61.5%, which was consistent with other species of musk deer (Table 3). The estimated average values of AT and GC skews in rRNA was found to be 0.211 and −0.098 receptively (Table 2). 22 tRNA genes were distributed with variable lengths from tRNA-Ser (60 bp) to (tRNA-Leu (75 bp). The overall AT and GC content in 1511 bp long tRNA was 64.5% and 35.5% respectively. The resultant value of AT and GC skew was 0.026 and 0.086, receptively (Table 2). Out of the 22 tRNAs, fourteen genes were located on H-strand while the other eight were on L-strand (Table 1).

**Table 3:** Nucleotide composition indices in different regions of mitogenomes of *Moschus* species.

| Species | Accession number | Whole Mitogenome |  | Protein Coding Genes (PCGs) |  | Ribosomal RNA (rrnl) |  | Control region |  |
| --- | --- | --- | --- | --- | --- | --- | --- | --- | --- |
|  |  | Length (bp) | AT (%) | Length (bp) | AT (%) | Length (bp) | AT (%) | Length (bp) | AT (%) |
| *M. cupreus* | MTXXXXXX | 16354 | 62.5 | 11404.0 | 62.3 | 2527.0 | 61.9 | 925 | 64.5 |
| *M. chrysogaster* | KP684123 | 16354 | 61.9 | 11404.0 | 61.6 | 2528.0 | 61.5 | 924 | 63.2 |
| *M. leucogaster* | NC042604 | 16431 | 62.0 | 11404.0 | 61.6 | 2528.0 | 61.6 | 1001 | 63.9 |
| *M. anhuiensis* | KP684124 | 16352 | 62.0 | 11404.0 | 61.8 | 2527.0 | 61.2 | 923 | 62.5 |
| *M. berezovskii* | EU043465 | 16354 | 62.1 | 11404.0 | 62.0 | 2528.0 | 61.2 | 924 | 62.8 |
| *M. moschiferus* | FJ469675 | 16351 | 62.2 | 11401.0 | 61.9 | 2528.0 | 61.5 | 923 | 63.7 |

### Mitochondrial control region sequence

The CR region was positioned between tRNAPro and tRNAPhe which was 925 bp long. The AT content was higher than the GC content with the base composition of CR region as 32.2% A, 32.3% T, 13.8% G and 21.6% C. The skew value of AT and GC for CR was 0.002 and −0.220, respectively (Table 2). The CR sequence of *M. cupreus* was found to be smaller as compared to the previously reported Himalayan Musk deer (*M. leucogaster*) sequence of 1001 bp with a 77 bp long insertion (Guo et al., 2018). The insertions and deletions (INDEL) have been studied in many mammalian species such as *Rucervus eldii:* 78 bp (Balakrishnan et al., 2003), *Homo sapiens:* 9 bp (Thangaraj et al., 2005), *Rusa unicolor:* 40 bp (Gupta et al., 2015), and *Rucervus duvaucelii:* 94 bp (Kumar et al., 2017).

### Phylogenetic relationship and genetic distance

The phylogenetic relationship was estimated based on the complete mitogenomes of *Moschus* species with other species of Cervini, Bovini, and Boselaphini. The result indicated all *Moschus* species clustering in a monophyletic clade distinct from other species of Cervini, Bovini, and Boselaphini (Fig. 3). Previously, *M. moschiferus* was reported as the most primitive species in genus *Moschus* (Vislobokova et al., 2009) while the study by Pan et al., 2015 suggested that the ancestral lineage of the present musk deer was likely to be distributed in the Tibet Plateau margin or adjacent mountains around the Sichuan Basin. Our phylogenetic tree indicated that *M. cupreus* was the basal clade in *Moschus*. This indicated *M. cupreus* to be the most primitive ancestral species in the genus *Moschus*.

**Figure 3:**
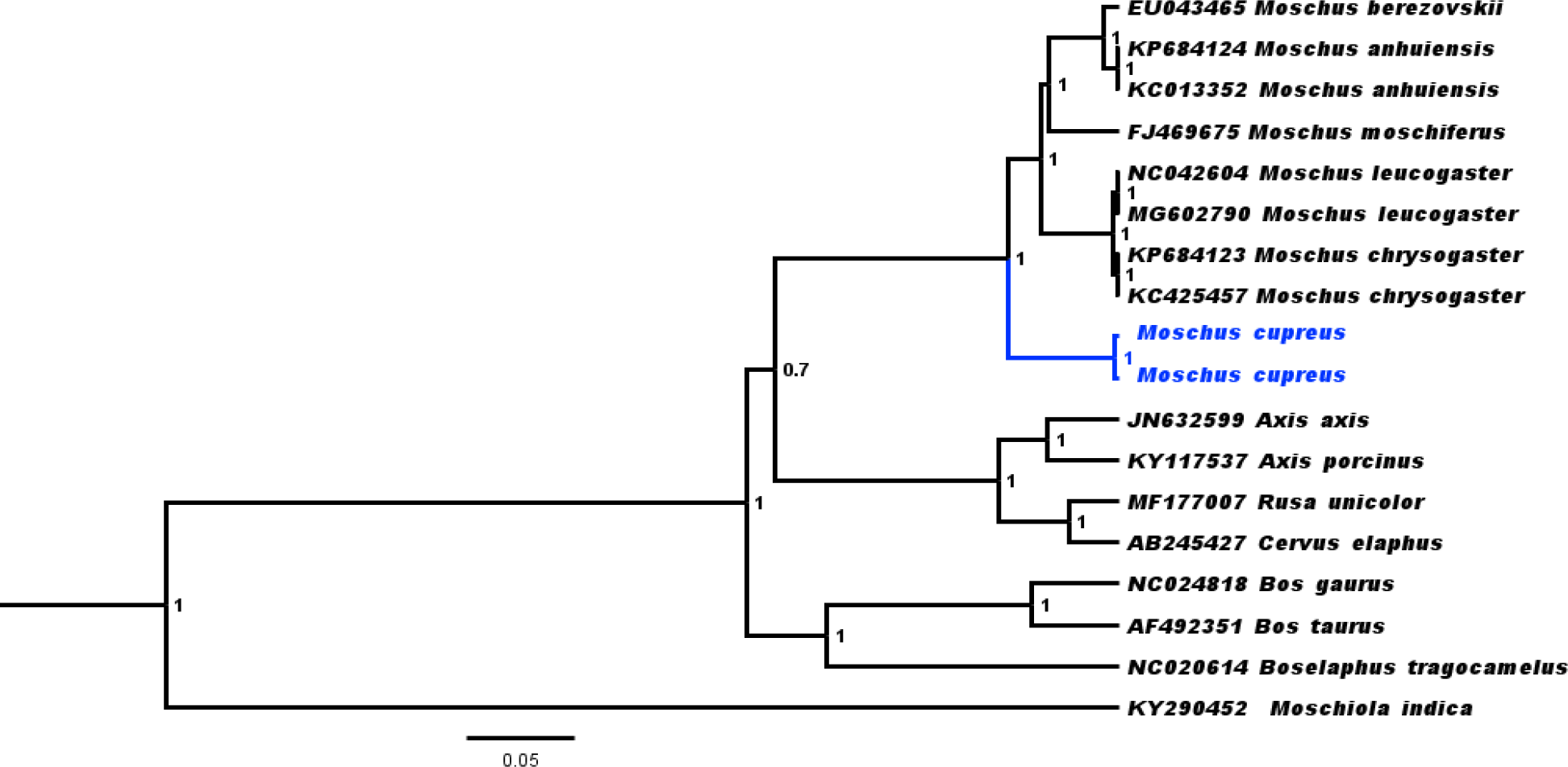
Phylogenetic relationship among musk deer and related species inferred from complete mitogenome using Bayesian inference (BI). Bayesian posterior probability values are shown at each node. *Moschiola indica* (KY290452) was used as an outgroup.

We calculated the genetic relatedness of *M. cupreus* with other musk deer species based on 13 PCGs, 2 rRNA and Control region (Fig. 4). The overall genetic similarity of *M. cupreus* indicated high genetic similarity (average 92%) with other musk deer species. We also calculated gene-wise genetic differentiation where results indicated varied genetic differentiation in all 13 PCG genes of musk deer (Fig. 5).

**Figure 4:**
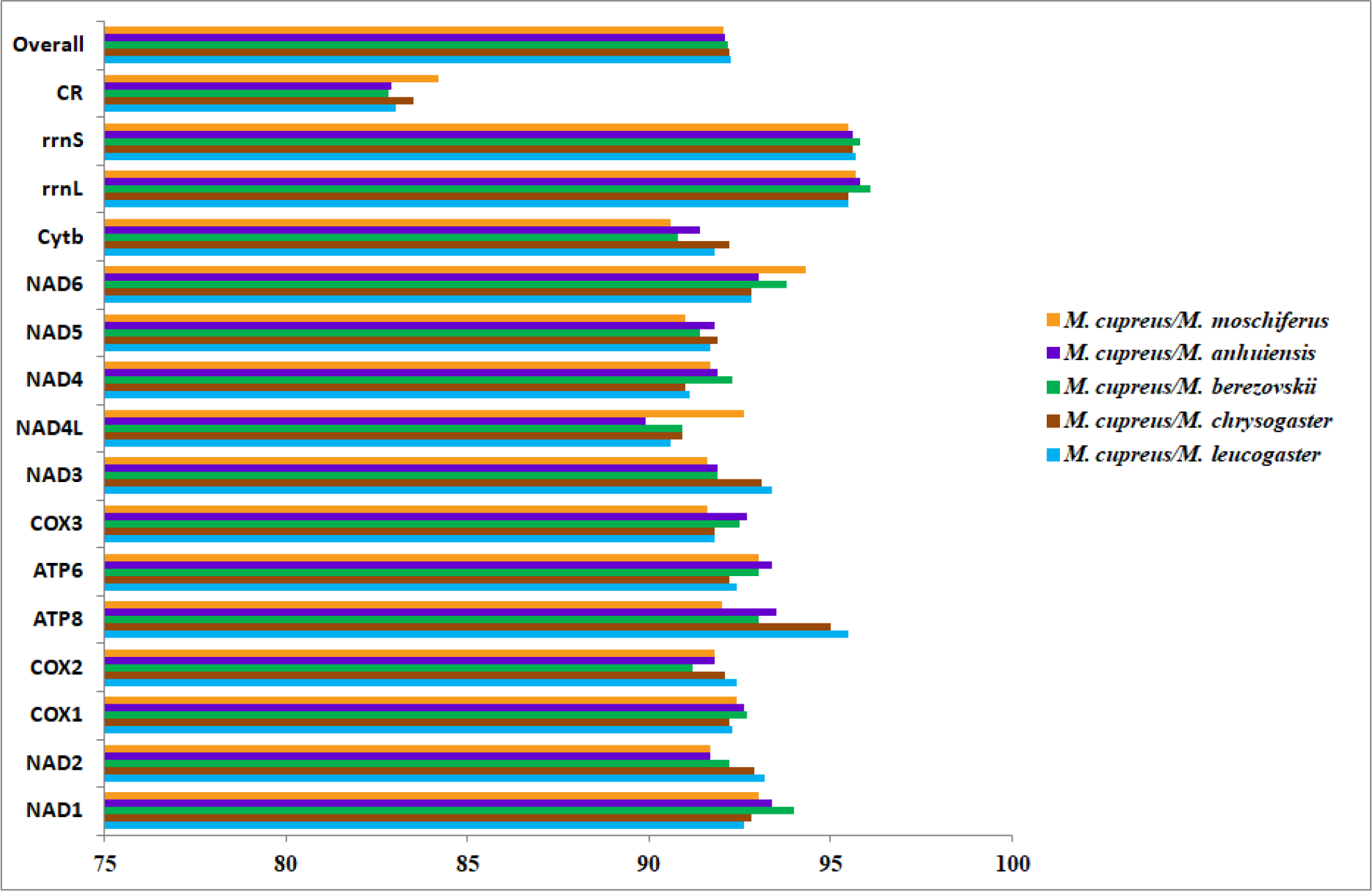
Graph showing the similarity of *M. cupreus* with other musk deer species.

**Figure 5:**
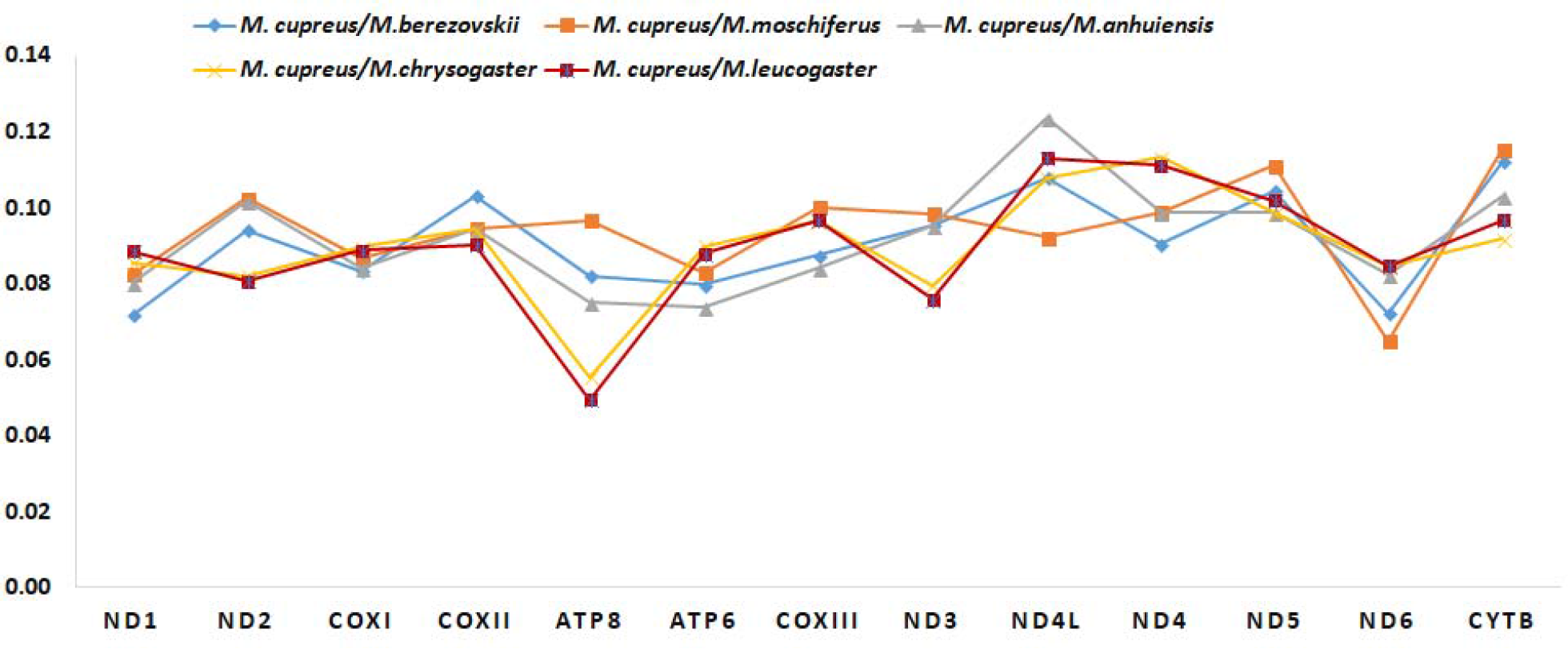
Comparative pairwise genetic distance in the protein-coding gene of musk deer species

## Conclusion

Previously, *M. cupreus* has been described as a subspecies of *M. chrysogaster* (Timmins et al., 2015). Based on morphological similarity, *M. cupreus* was also confused with *M. leucogaster*. Thereafter, Groves et al., 1995 speculated that it might be a separate species, which was subsequently confirmed by Grubb 2005. Evolutionary history and phylogenetic position of *M. cupreus* in the genus *Moschus* was still uncertain due to limited genetic information. The novel complete mitogenome of Kashmir Musk deer (*M. cupreus*) will aid in accurate species identification and form baseline information for further exploration into *Moschus* and related species. Availability of large scale mitogenomic data will also help enforcement agencies in molecular tracking of seized wildlife items from musk deer apart from accurately guiding in-situ and ex-situ management strategies to efficiently conserve musk deer. It will also help in wildlife forensics for the identification of musk deer body parts (musk pod, canine and meat) recovered in the illegal wildlife trade.

## Acknowledgment

We thank Dr. Dhananjai Mohan, Director, Dr. Y.V. Jhala, Dean, and Dr. G.S. Rawat (former Dean and Director), WII for their support. We thank the State Forest Departments of Jammu and Kashmir and Uttarakhand for forwarding biological samples for the forensic examination to Wildlife Forensic and Conservation Genetics (WFCG) Cell, WII. We acknowledge the support provided by Mr. A. Madhanraj, WFCGC during this study.

